# Distinct components of photoperiodic light are differentially encoded by the mammalian circadian clock

**DOI:** 10.1101/2020.01.14.906131

**Authors:** Michael C. Tackenberg, Jacob J. Hughey, Douglas G. McMahon

## Abstract

Seasonal light cycles influence multiple physiological functions and are mediated through photoperiodic encoding by the circadian system. Despite our knowledge of the strong connection between seasonal light input and downstream circadian changes, less is known about the specific components of seasonal light cycles that are encoded and induce persistent changes in the circadian system. Using combinations of three T cycles (23, 24, 26 hr.) and two photoperiods per T cycle (Long and Short, with duty cycles scaled to each T cycle), we investigate after-effects of entrainment to these six light cycles. We measure locomotor behavior duration (α), period (τ), and entrained phase angle (Ψ) *in vivo*, and SCN phase distribution (σ_ϕ_), τ, and Ψ *ex vivo* in order to refine our understanding of critical light components for influencing particular circadian properties. We find that photoperiod and T cycle length both drive determination of *in vivo* Ψ but differentially influence after-effects in α and τ, with photoperiod driving changes in α and photoperiod length and T cycle length combining to influence τ. Using skeleton photoperiods, we demonstrate that *in vivo* Ψ is determined by both parametric and non-parametric components, while changes in α are driven non-parametrically. Within the *ex vivo* SCN, we find that Ψ and σ_ϕ_ of the PER2∷LUCIFERASE rhythm follow closely with their likely behavioral counterparts (Ψ and α of the locomotor activity rhythm), while also confirming previous reports of τ after-effects of gene expression rhythms showing negative correlations with behavioral τ after-effects in response to T cycles. We demonstrate that within-SCN σ_ϕ_ changes, thought to underly α changes *in vivo*, are induced primarily non-parametrically. Taken together, our results demonstrate distinct components of seasonal light input differentially influence Ψ, α, and τ, and suggest the possibility of separate mechanisms driving the persistent changes in circadian behaviors mediated by seasonal light.

## INTRODUCTION

The duration of daylight, or photoperiod, represents a predictable and dynamic signal of seasonal change throughout the year across much of the planet. The predictability of this signal is harnessed by organisms to initiate or avoid a variety of biological functions at specific times of year, including reproductive behaviors (Elliott and Goldman, 1981), resource conservation (Bartness et al., 1989; Bartness and Wade, 1984), and migration (Gwinner, 1990). In humans, a variety of disorders and non-communicable diseases have seasonal correlations (Basnet et al., 2016; Oh et al., 2010; Wehr, 2001), with perhaps the most striking seasonal influence being on depression-anxiety behaviors (Koorengevel et al., 2003). Despite clear associations between photoperiodic exposure and physiological function and dysfunction, the roles of specific components of seasonal light cycles that induce those changes are incompletely understood.

One factor critical to our current understanding of the mechanism of photoperiodic induction is the persistence of seasonal changes. Physiological hallmarks of exposure to particular light schedules can be observed, for varying intervals, beyond direct exposure to that light schedule (Pittendrigh and Daan, 1976a). These persistent changes, referred to as after-effects, include characteristic changes to core circadian locomotor behavior patterns in the form of activity duration (alpha, α), free-running period (tau, τ), and phase angle of entrainment (psi, Ψ). In nocturnal animals, prior exposure to long photoperiods results in after-effects of compressed α, a shortened τ, and an advanced Ψ relative to short photoperiod counterparts as measured upon release into DD (Pittendrigh and Daan, 1976a).

The existence of after-effects suggests that seasonal light cycle information is encoded within the brain and can persist beyond the interval of direct exposure. The master circadian pacemaker of the brain, the suprachiasmatic nucleus (SCN), likely plays a major role in this encoding, particularly with regards to after-effects in circadian behavior (for review, see Tackenberg and McMahon, 2018). In addition, recent work has identified a distinct non-SCN target for photoperiodic influence, the peri-habenular area, influencing seasonal response in mood (Fernandez et al., 2018). Like circadian locomotor behavior patterning, the SCN undergoes substantial changes in organization following exposure to different photoperiods, constituting network-level after-effects. These responses include changes in phase distribution (σ_ϕ_) of its constituent neurons as well as in τ and Ψ of SCN gene expression rhythms. Prior exposure to long photoperiod has been shown to increase σ_ϕ_ of *Period1∷Luciferase* (Inagaki et al., 2007), PERIOD2∷LUCIFERASE (Buijink et al., 2016), and electrical (VanderLeest et al., 2007) rhythms compared to short photoperiod counterparts. As in locomotor behavior after-effects, some reports have shown that prior exposure to long photoperiod shortens the SCN τ compared to short photoperiod counterparts (16:8 L:D, Ciarleglio et al., 2011; 20:4 L:D, Evans et al., 2013) while others have found non-significant decreases in SCN τ length following exposure to long photoperiod, particularly in the anterior SCN (Buijink et al., 2016; Mickman et al., 2008).

Induction of persistent circadian after-effects is not limited to changes in photoperiod. Manipulation of total day-night cycle (T cycle) length also induces after-effects in circadian locomotor behavior, with entrainment to short T cycles (T < 24 hours) resulting in contracted α (Azzi et al., 2014) and shortened τ upon release into constant conditions (Azzi et al., 2014; Schwartz et al., 2011). However, unlike photoperiodic after-effects, in which the changes in the τ of the explanted SCN itself match those on behavior, there is a negative correlation between behavioral and SCN explant τ following entrainment to T cycles. Exposure to short T cycles results in a long SCN τ and exposure to long T cycles results in a short SCN τ (Aton et al., 2004; Azzi et al., 2017; Molyneux et al., 2008).

The findings described above reveal a potential pattern of circadian after-effects, with long photoperiods and short T cycles producing one set of behavioral responses (short α, short τ), and short photoperiods and long T cycles producing another (long α, long τ). Because recent work has revealed a role for DNA methylation in establishing τ after-effects of extended T cycle entrainment, the potential association between τ and α after-effects provides a promising biochemical mechanism for both changes (Azzi et al., 2014). The linkage between the two types of after-effects, however, has not yet been sufficiently interrogated. The discordance between the SCN after-effects of T cycle and photoperiod offers a hint that the relationship may not be so simple. Here, we seek to further examine the alignment of T cycle and photoperiodic after-effects by measuring *in vivo* (α, τ, and Ψ of locomotor behavior rhythms) and *ex vivo* (σ_ϕ_, τ, and Ψ of SCN gene expression rhythms) responses following exposure to six combinations of photoperiod length and T cycle. We find that after-effects in each of these rhythm characteristics respond to the six different light cycles in distinct patterns, suggesting that they are each induced by different aspects of the input light cycle. Using skeleton photoperiods, we demonstrate that parametric vs. non-parametric responses differ between after-effects in α, τ and Ψ, further indicating encoding of distinct components of light input by each of these characteristics. Examining the rhythms of explanted SCN *ex vivo* following entrainment to these light schedules, we find that the pattern of SCN σ_ϕ_ reflects that of α *in vivo*, providing additional correlative evidence that SCN σ_ϕ_ may be a determining factor in setting α.

## MATERIALS AND METHODS

### Animals

Animal experiments were conducted in accordance with Vanderbilt University Institutional Animal Care and Use Committee regulations. All animals used for behavioral experiments were heterozygous for the *Per2∷Luciferase* allele (*Per2∷Luciferase*^*-/-*^) or were *Per2* wild-type (*Per2∷Luciferase*^*-/-*^). All animals used for SCN slice culture experiments were heterozygous for the *Per2∷Luciferase* allele (*Per2∷Luciferase*^-/-^).

### Activity Monitoring and Housing

Animals were singly-housed in light-tight boxes with activity monitoring by wheel revolutions detected by ClockLab acquisition software (Actimetrics, Inc.). Complete photoperiods used were 15:8 and 16:7 (T23 Long, ∼67% duty cycle), 8:15 and 7:16 (T23 Short, ∼33% duty cycle), 16:8 (T24 Long, ∼67% cycle), 8:16 (T24 Short, ∼33% cycle), 17:9 (T26 Long, ∼67% cycle), and 9:17 (T26 Short, ∼33 cycle). Animal numbers by group were as follows: 15:8 (*in* vivo n = 5, *ex vivo* n = 4), 16:7 (5, 2), 8:15 (6, 4), 7:16 (2, 0), 16:8 (12, 6), 8:16 (12, 8), 17:9 (6, 7), 9:17 (14,6). The two photoperiod duty cycles used in T23 (15:8/8:15 vs. 16:7/7:16) gave similar results and were combined into T23 Long and T23 Short groups for analysis.

Animals were transferred to the specified light schedule after at least 3 weeks of age, and housed there for at least 28 days before transfer to DD or brain extraction for slicing. Six T26 Short animals included in the datasets (three used in behavioral experiments, three used in slice experiments) experienced a multi-hour light failure during the 28 days of entrainment, but received 14 days of the correct light cycle following the failure.

Skeleton photoperiods were set up by housing animals in 12:12 LD complete photoperiods for approximately 5 days before transitioning to a 12:12 skeleton (1:10:1:12 L:D:L:D). After several cycles on this skeleton, the onset pulse was gradually advanced (0.5 hour – 1 hour per day) or delayed (1 hour per day) until a skeleton long (1:14:1:8 L:D:L:D) or skeleton short (1:6:1:16 L:D:L:D) photoperiod, respectively, was established. Once the final skeleton was established, animals were given 28 days of exposure to that skeleton until transfer to DD or slicing. Animal numbers by group were as follows: skeleton short (*in vivo* n = 6, *ex vivo* n = 5), skeleton long (7, 6).

### Actogram Analysis

Locomotor behavior was analyzed using ClockLab Analysis software and R. Period was determined using the Chi Square periodogram over the first 7 days of DD. α was measured using automated onset/offset detection for each cycle of the actogram. Full actograms can be found in Supplemental Data 1.

### Automated α Measurement

Automated α measurement was performed in R. The activity (bin size 6 minutes) is smoothed by a Savitzky-Golay filter with a span of 25 and degree of 3. A threshold is set relative to the maximum smoothed activity (0.01 * the maximum for each day). Points at which the smoothed activity crosses this threshold are identified as potential onsets and offsets. Segments of activity less than 2 hours are considered inactive. The longest inactive segment is identified, with the start of that segment recorded as the offset and the end of that segment recorded as the onset. α is calculated as activity offset – onset, and Ψ as the timing of the activity onset in the first cycle in DD relative to the projected time of lights off from the previous light dark cycle (projected Zeitgeber Time 12 [ZT12]). Identified onsets and offsets for each cycle can be found in Supplemental Data 2. R scripts used for analysis can be found at 10.6084/m9.figshare.11513892.

### Slicing and Slice Cultures

Within 4 hours of lights off, animals were sacrificed by cervical dislocation and the brain removed. SCN slices of 200-μm were made, and the SCN further isolated by scalpel cut under a dissecting microscope. Slices were transferred to a 6-well plate with cell culture membrane insert (Millipore, PICMORG50) and 1.2-mL of slice culture media. Slice chambers were sealed with a coverslip and vacuum grease and placed into an incubated light-tight inverted microscope (Zeiss Axioskop). Luminescence was detected using an intensified CCD (Stanford Photonics) controlled by μ- Manager recorded for 2’ every 10’ for at least 1 week.

### *Ex-Vivo* Analyses

Luminescence recordings were analyzed using Fiji and R. OME-TIFF files from μ-Manager were opened in Fiji and smoothed using a two-frame minimization (frame rate reduction from 6/hour to 3/hour). A 102×102 grid of 10×10-pixel (5.175 × 5.175 μm) ROIs was overlaid on the images and brightness measured for each of these 10,404 ROIs measured for every frame. These grid measurements were then exported to R for further analysis. The first 8 hours of recording were excluded from peak-finding to prevent slicing artifacts from interfering with measurements. First, the ROIs comprising the SCN were identified by using pixels within the top 50% of brightness. The timing of each peak of luminescence was then detected for each ROI and used to calculate the initial phase. Distribution measurements were made by median absolute deviation across all SCN ROIs. Phase maps of each SCN used can be found in Supplemental Data 3. R scripts used for analysis can be found at 10.6084/m9.figshare.11513892. Cycle-by-cycle interpeak-calculated period and phase distribution are shown in Supplementary Figure 1 and 2, respectively.

**Figure 1.**
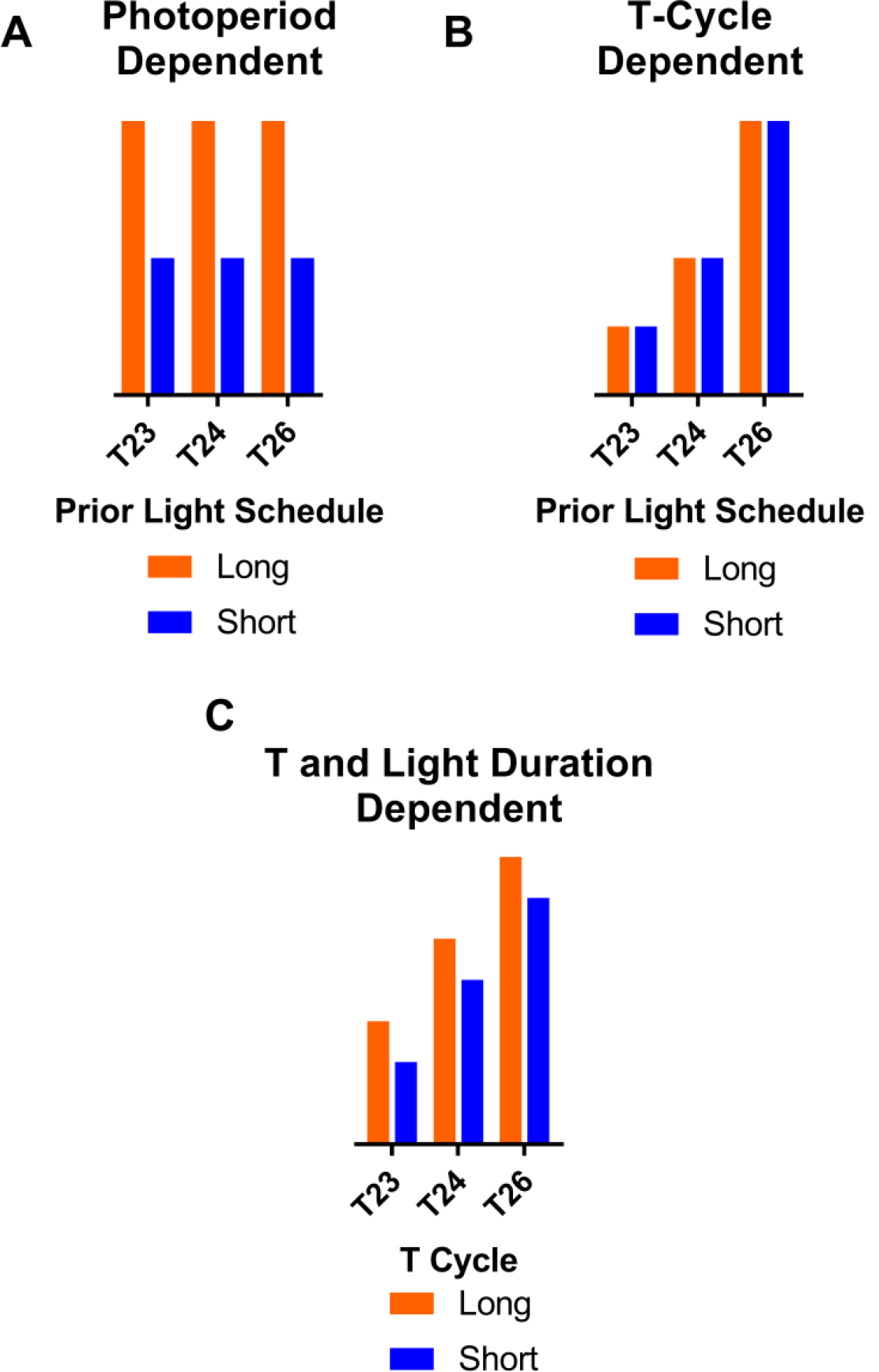
Schematic of possible patterns of measurements across T23 L/S, T24 L/S, and T26 L/S. **A**, a T-cycle dependent response where the measurement is graded depending on the T cycle and photoperiod within the T-cycle has negligible effects. **B**, a photoperiod-dependent response where light duty cycle dictates the measurement regardless of T cycle. **C**, a combination of photoperiod and T cycle influences, with each of the 6 combinations having a characteristic measurement level. Note that in all examples, the polarity of the measurement change is arbitrary (e.g., T cycle response in A can increase or decrease with T cycle, photoperiod response in B can have either Long or Short measure higher).These patterns are illustrations of individual and combined main effects; possible interaction effects are not shown.

**Figure 2.**
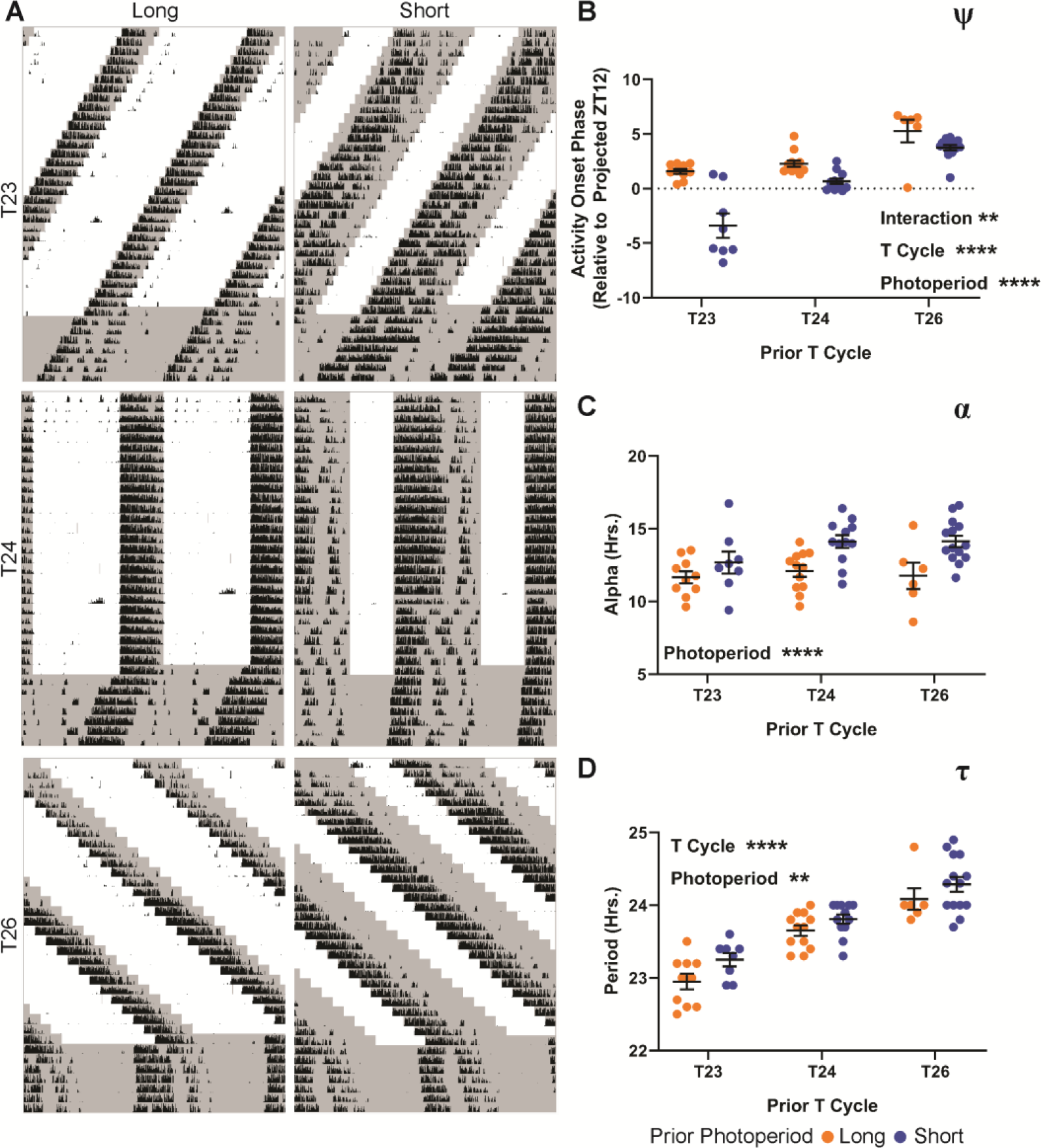
Behavioral after-effects of six photoperiod/T cycle combinations. **A**, representative actograms from long (left column) and short (right column) photoperiods combined with T23 (top row), T24 (middle row), and T26 (bottom row). **B**, Ψ following exposure to the six light schedules. Values are relative to projected ZT12 (positive values advanced, negative values delayed). Interaction *p* = 0.0014 (7.242%), T Cycle Length *p* < 0.0001 (51.30%), Photoperiod Legth *p* < 0.0001 (20.97%). **C**, α after-effect across all six schedules as measured by automated α detection (see Methods). Interaction *p* = 0.4256 (2.081%), T Cycle Length *p* = 0.1648 (4.468%), Photoperiod Length *p* < 0.0001 (21.67%). **D**, period after-effect across the six schedules as measured by chi square periodogram. Interaction *p* = 0.7587 (0.2835%), T Cycle Length *p* < 0.0001 (56.17%), Photoperiod Length *p* = 0.0087 (3.778%). Two-way ANOVA results represent *p* value and percent variation (*p* < 0.0001 ****, *p* < 0.001 ***, *p* < 0.01 **, *p* < 0.05 *). T23 Long, n = 10; T23 Short, n = 8; T24 Long, n = 12; T24 Short, n = 12; T26 Long, n = 6; T26 Short, n = 14. T23 Long and Short groups are composed of a combination of two similar LD ratios (see Methods).

### Statistical Analysis

For measurements of α, τ, σ_ϕ_, and Ψ, the effects of photoperiod and T cycle (or completeness and onset/offset interval) were analyzed by two-way ANOVA. For each comparison, the *p* value and percent of total variation of each factor, as well as their interaction, are reported. Where indicated in the text, Sidak’s multiple comparisons test was used to further examine within- and between-factor effects. For correlation analysis shown in Figure 3, Pearson’s r test was used.

**Figure 3.**
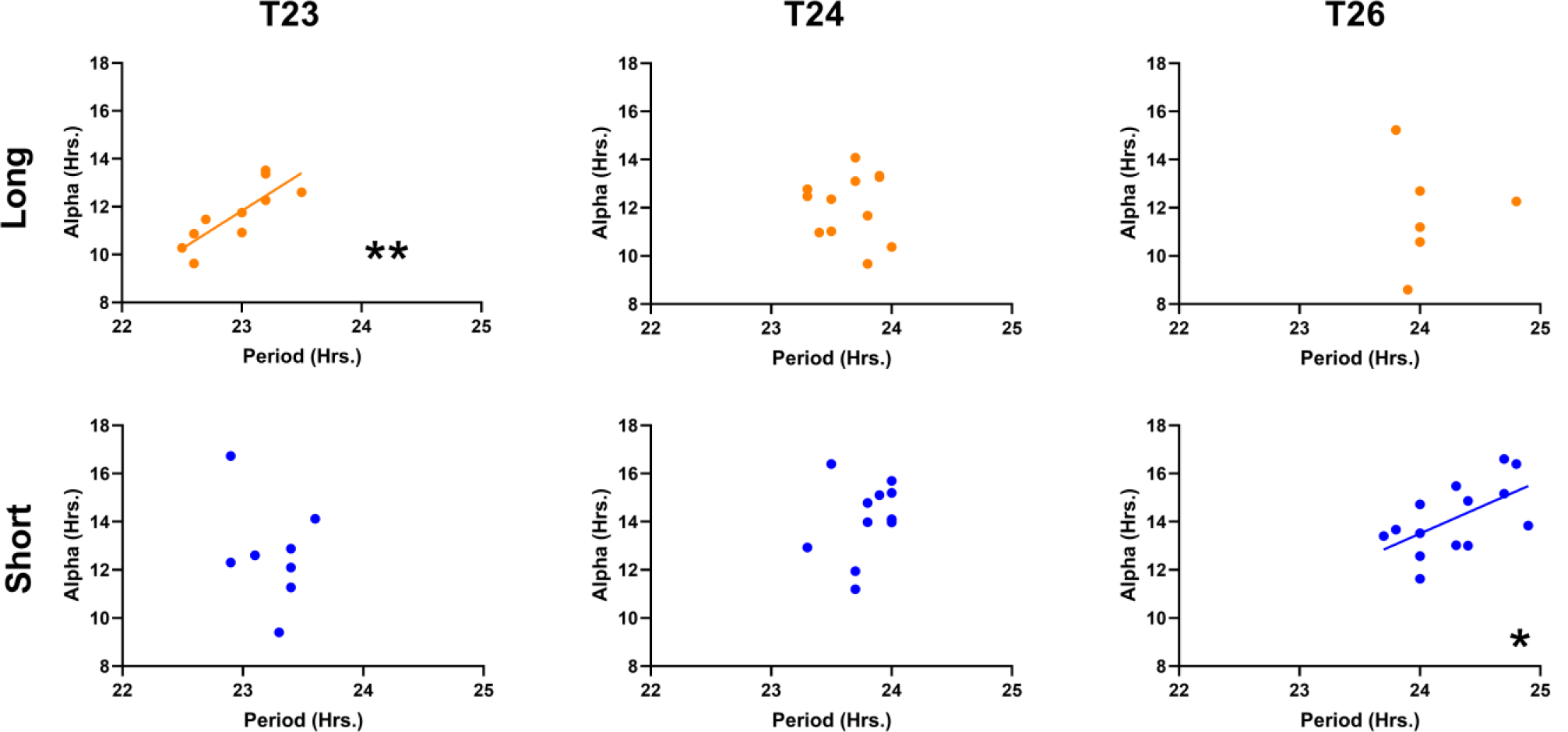
Within-individual correlations between τ and α after-effects following the given T-cycle/photoperiod combination. T23 Long 0.8238 (*p* = 0.0034, **), T24 Long −0.06835 (*p* = 0.8328, ns), T26 Long 0.004966 (*p* = 0.9926, ns), T23 Short −0.3147 (*p* = 0.4477, ns), T24 Short 0.2671 (*p* = 0.4013, ns), T26 Short 0.5818 (*p* = 0.0291, *); Values reported are Pearson r and *p* value (*p* < 0.0001 ****, *p* < 0.001 ***, *p* < 0.01 **, *p* < 0.05 *).

## RESULTS

### Distinct response patterns of Ψ, and after-effects on α and τ

To examine the persistent encoding of photoperiods and T cycles *in vivo*, as represented by entrained phase angle (Ψ) and after-effects in activity duration (α) and free-running period (τ), we exposed mice to six different light schedules consisting of combinations of short or long photoperiod and short, standard, and long T cycles (T23 Long, T23 Short, T24 Long, T24 Short, T26 Long, T26 Short). Following 14-28 days of exposure to these light schedules (see Methods), animals were transferred into constant darkness (DD) for 7 days. Based on previous studies examining the after-effects of T cycles (Azzi et al., 2014, 2017; Pittendrigh and Daan, 1976a; Schwartz et al., 2011) and photoperiod (Buijink et al., 2016; Ciarleglio et al., 2009; Evans et al., 2013; Pittendrigh and Daan, 1976a), we hypothesized that the response of τ and α after-effects to these six light schedules would produce one of three general patterns (Figure 1).

First, if after-effects in these rhythm characteristics were strictly dependent on the duty cycle of the light interval, then the groups would segregate into two levels, corresponding to long or short photoperiods, regardless of T cycle (Figure 1A). A primary influence of the photoperiod could indicate a parametric effect of the total light duration per cycle and/or a non-parametric effect from the phase shifts caused by light onset and light offset. Second, if plasticity is strictly dependent on net daily phase advances or delays driven by entrainment to the T cycles, then the measurements would segregate into three groups based on T cycle (Figure 1B). This pattern of results would most readily suggest that after-effects were induced by non-parametric effects of the daily entraining phase shifts induced by the T cycles. Third, significant main effects of both T cycle and photoperiod would result in after-effects with each of the 6 groups having a characteristic level (Figure 1C), while interactions between T cycles and photoperiod would produce many possible variations of the pattern in Figure 1C.

Indeed, Ψ of locomotor rhythms relative to projected ZT12 of the previous light:dark cycle (see Methods) was strongly influenced by both T cycle length and photoperiod length, with a significant interaction between these two main effects (Figure 2B). As T cycle length increases, Ψ becomes more advanced. Within each T cycle, the short photoperiod version produces a more delayed Ψ than its long photoperiod counterpart, but the difference between the two photoperiods decreases as T cycle length increases. This pattern resembled the example in Figure 1C, with the interaction providing an additional layer of complexity. In contrast to the multifaceted inputs to Ψ, we found that photoperiod was the sole significant influence on α, with persistently reduced α in DD following long photoperiods compared to short photoperiods across all three T cycles (Figure 2C). This pattern most closely resembled the example in Figure 1A, and suggested that a parametric effect of the duration of light per cycle and/or a non-parametric effect of the phase shifts caused at light onset and offset induced the after-effect in α. After-effects on locomotor τ were significantly influenced by both T cycle length and photoperiod (Figure 2D), but unlike Ψ, there was no detectable interaction between the main effects. Within each T cycle, the long photoperiod group had consistently shorter τ compared to its partner short photoperiod. Across T cycles, however, there was a direct relationship with τ, with τ increasing along with T cycle length regardless of photoperiod. This pattern most closely resembles the example in Figure 1C.

To further examine the relationship between the α and τ after-effects, we plotted the two measurements against one another (Figure 3). The two factors correlated significantly in two specific cases: T23 Long (Figure 3, top left) and T26 Short (Figure 3, bottom right). These two photoperiod/T cycle combinations represent the instances where in previous reports T cycle and photoperiod independently produce similar *in vivo* after-effects on α and τ, with long photoperiod or short T decreasing α and τ, and short photoperiod or long T increasing α and τ. Interestingly, there was a lack of correlation of these measures in T cycle/photoperiod combinations where the predicted after-effects of the cycle τ and photoperiod would be in nominal conflict (Long photoperiod and Long T, and Short photoperiod and Short T, Azzi et al., 2014; Pittendrigh and Daan, 1976a).

### Ψ and α after-effects differ in their parametric vs. non-parametric responses

The after-effects described in Figure 2 for all parameters showed significant main effects of photoperiod, revealing that either the interval between light transitions or the light duration is inducing persistent effects on Ψ, α, and τ. To determine whether the critical aspect of the light cycle is the timing of the light/dark and dark/light transitions or the duration of light within each cycle, we used skeleton photoperiods consisting of two 1-hour pulses of light, separated by intervals of darkness, that mimic the timing of the light/dark transitions on full photoperiods of 16:8 and 8:16 (Figure 4A). Skeleton photoperiods have been used to assess the mechanism of entrainment through seasonal changes in Ψ (Pittendrigh and Minis, 1964), as well as to determine the proximal effect on locomotor behavior duration (Pittendrigh and Daan, 1976a, 1976b). In order to determine whether skeleton photoperiods can induce plasticity in α, τ, and Ψ, we assessed after-effects in locomotor behavioral rhythms in DD following 28 days of exposure to skeleton long or short photoperiods in comparison to similar entrainment to full 16:8 and 8:16 photoperiods.

**Figure 4.**
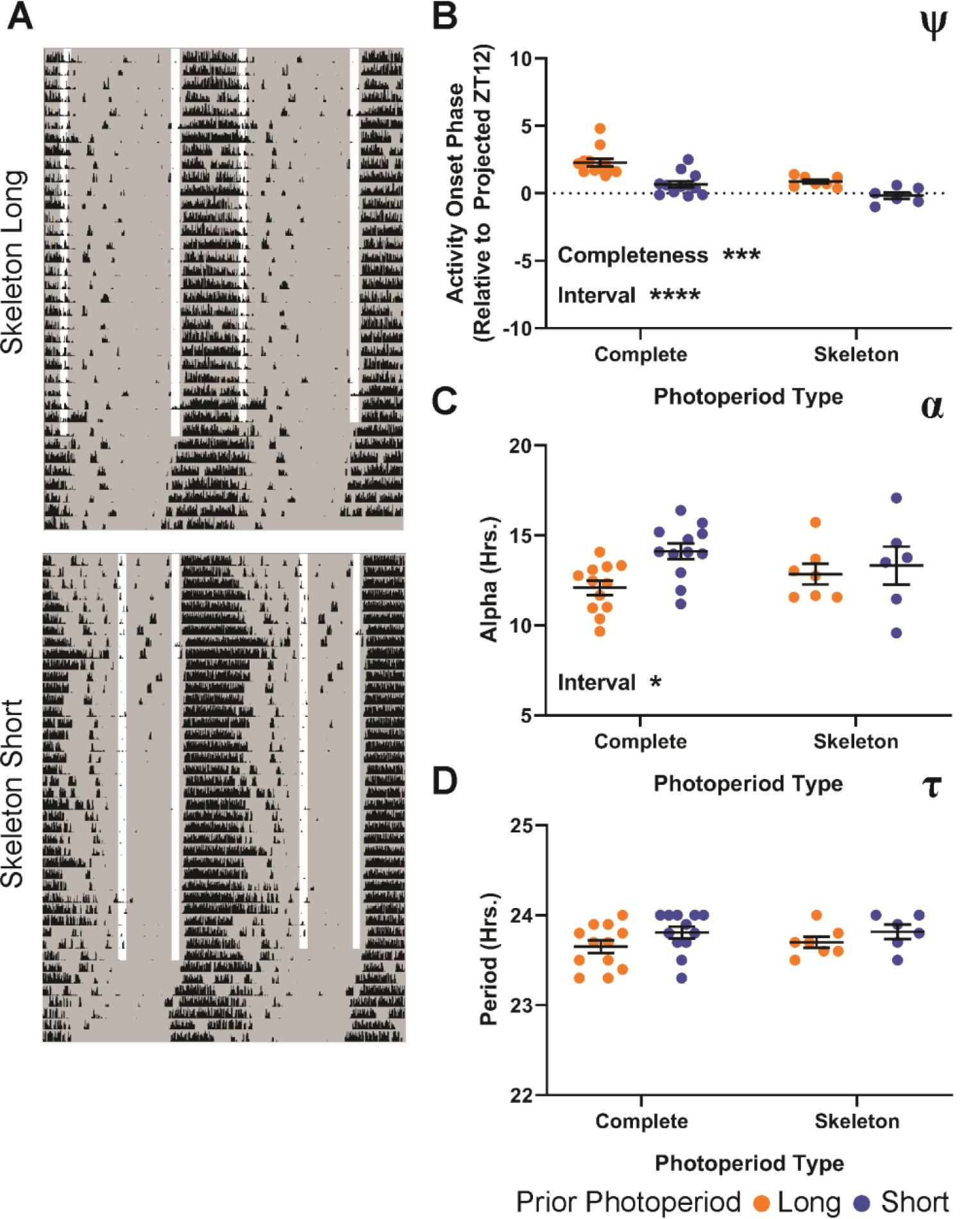
Behavioral after-effects following exposure to skeleton long and short photoperiods compared to T24 complete long and short photoperiods. **A**, representative actograms from long (top) and short (bottom) skeleton photoperiods. **B**, Ψ following exposure to skeleton long and short photoperiods. Values are relative to projected ZT12 (positive values advanced, negative values delayed). Interaction *p* = 0.3161 (1.33%), Completeness *p* = 0.0003 (21.24%), Onset-Offset Interval *p* < 0.0001 (29.32%). **C**, α after-effect of skeleton long and short photoperiods. Interaction *p* = 0.1861 (4.332%), Completeness *p* = 0.9858 (0.0008%), Onset-Offset Interval *p* = 0.0376 (11.15%). **D**, τ after-effect of skeleton long and short photoperiods. Interaction *p* = 0.7835 (0.2063%), Completeness *p* = 0.7006 (0.4044%), Onset-Offset Interval *p* = 0.0765 (8.988%). Long and Short complete photoperiod values in B-D are replotted from Figure 2. Two-way ANOVA results represent *p* value and percent variation (*p* < 0.0001 ****, *p* < 0.001 ***, *p* < 0.01 **, *p* < 0.05 *). Complete Long, n = 12; Complete Short, n = 12; Skeleton Long, n = 7, Skeleton Short, n = 6.

Ψ exhibited main effects of both the duration of light in complete photoperiods (Completeness, Figure 4B), and the timing of light transitions in skeleton photoperiods (Interval, Figure 4B). The significant effect of the light duration of complete photoperiods indicates that there is a parametric influence on the Ψ of locomotor behavioral rhythms, in addition to the non-parametric main effect of the interval between light transitions on skeleton photoperiods. In contrast, we found a significant main effect only of the onset-offset interval of skeleton photoperiods on α after-effects, suggesting that this effect is primarily non-parametric (Figure 4C). We found that the complete and skeleton photoperiods had no significant after-effect effect on τ (Figure 4D), however previous investigators have reported τ after-effects of complete and skeleton photoperiods (Pittendrigh and Daan, 1976a). Thus we found that there are significant parametric and non-parametric effects on Ψ, non-parametric effects of onset-offset interval on α, and non-significant after-effects on behavioral τ of both complete and skeleton photoperiods.

### *Ex vivo* SCN after-effects

Using *ex vivo* imaging of PER2∷LUCIFERASE (PER2∷LUC) rhythms following entrainment to the same six light schedules described above, we investigated the effects of photoperiod and T cycles on plasticity of the SCN pacemaker. We housed heterozygous PER2∷LUC animals in identical conditions to our complete photoperiod *in vivo* experiments, including running wheels, and collected SCN slices from the animals after 14-28 days of exposure to each of the light schedules described above. Slices were collected within 4 hours of lights off, maintained in organotypic culture, and PER2∷LUC luminescence recorded *ex vivo* for 5-7 days (see Methods). Slice recordings were analyzed by detecting peak PER2∷LUC expression in each of a 102 × 102 grid of sub-cellular (∼5 × 5 μm) regions of interest (ROIs) spread across the entire SCN image. These PER2∷LUC peak times were then used to calculate and map σ_ϕ_ (Figure 5A), while the average PER2∷LUC luminescence profile over time was used to calculate SCN τ.

**Figure 5.**
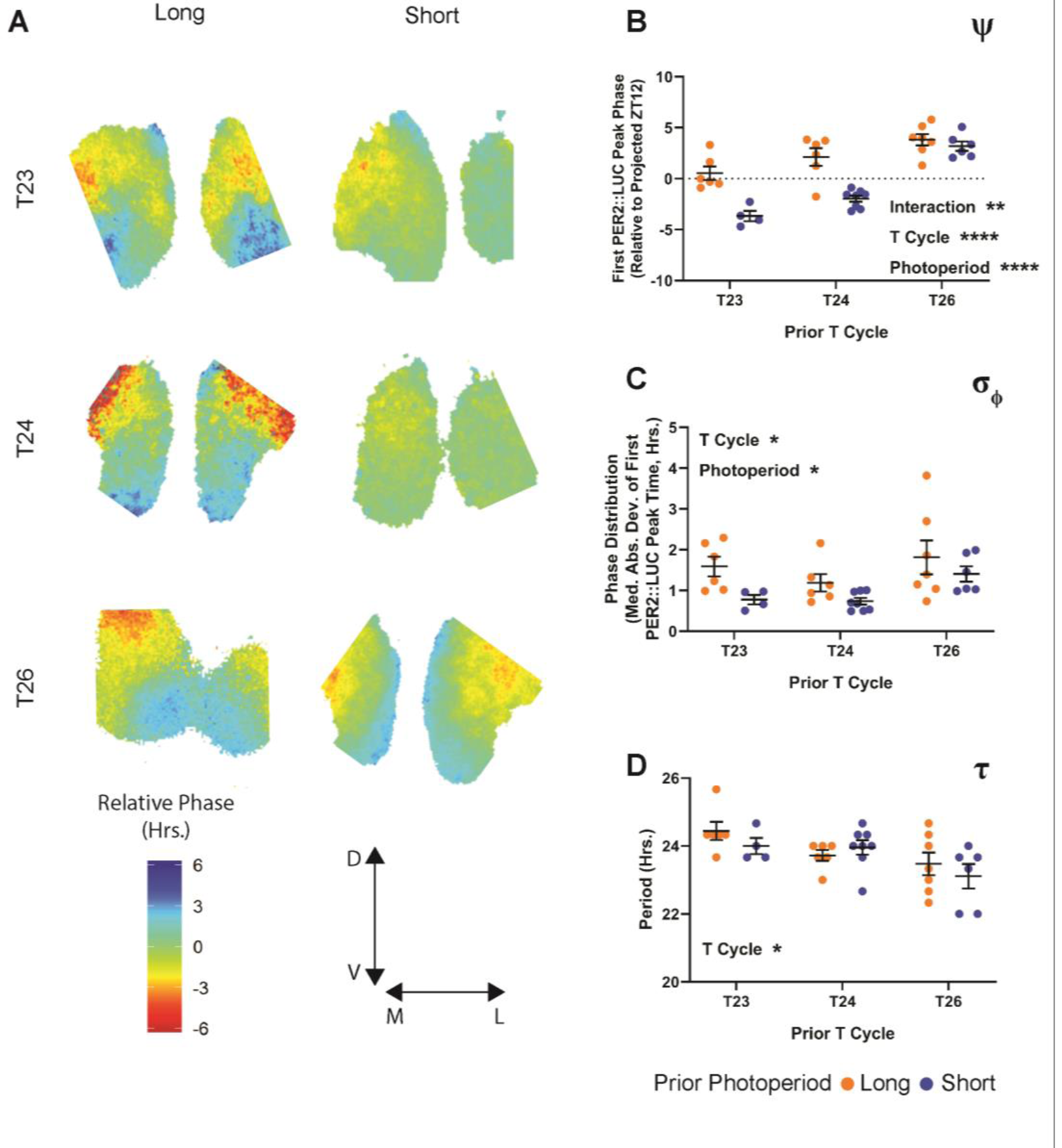
PER2∷LUC after-effects following exposure to six photoperiod/T cycle combinations. **A**, representative relative phase maps from long (left column) and short (right column) photoperiod combined with T23 (top row), T24 (middle row), or T26 (bottom row). **B**, Ψ of the PER2∷LUC rhythm following entrainment to the specified T cycle/photoperiod combination. Values represent the time of the first PER2∷LUC peak relative to the projected lights off for the first cultured cycle (projected ZT 12). Interaction *p* = 0.0052 (8.185%), T Cycle Length *p* < 0.0001 (49.47%), Photoperiod Length *p* < 0.0001 (24.94%). **C**, σ_ϕ_ (median absolute deviation of PER2∷LUC peak times) across SCN slices from animals entrained to each of the specified light schedules. Interaction *p* = 0.7168 (1.433%), T Cycle Length *p* = 0.0353 (15.89%), Photoperiod Length *p* = 0.0114 (15.42%). **D**, τ of the PER2∷LUC rhythm over 5 cycles *ex vivo*. Interaction *p* = 0.4104 (4.109%), T Cycle Length *p* = 0.0111 (23.40%), Photoperiod Length *p* = 0.4151 (1.529%). Two-way ANOVA results represent *p* value and percent variation (*p* < 0.0001 ****, *p* < 0.001 ***, *p* < 0.01 **, *p* < 0.05 *). T23 Long, n = 6; T23 Short, n = 4; T24 Long, n = 6; T24 Short, n = 8; T26 Long, n = 7; T26 Short, n = 6. T23 Long is composed of a combination of two similar LD ratios (see Methods).

Similar to the Ψ of locomotor rhythms, we found that the Ψ of SCN PER2∷LUC rhythms (relative to projected ZT12 of the previous light cycle) was significantly influenced by both T cycle and photoperiod with a significant interaction between the two factors (Figure 5B). As T cycle length increases, SCN Ψ also becomes more advanced, and within each T cycle the long photoperiod is more advanced than the short. Again, like the *in vivo* Ψ, the *ex vivo* Ψ featured a narrowing difference between long and short as T cycle length increases. Similarly, mirroring *in vivo* α, we found that the photoperiod modulated the degree of synchrony of PER2::LUC rhythms within explanted SCN (σ_ϕ_, Figure 5C), with increased phase dispersion in long photoperiods compared to short. However, we also found a significant main effect of T cycle length on the SCN σ_ϕ_ after-effect. This effect was driven by increased σ_ϕ_ in both T26 groups compared to both T24 groups (post-hoc multiple comparisons test between T cycle lengths, T24 vs. T26 adjusted *p* = 0.0329). There is no single consistent linear effect of T cycle on the *ex vivo* σ_ϕ_ after-effect. As such, the pattern most resembled that of the example in Supplemental Figure 2A, but with the T26 groups elevated.

SCN τ after-effects following T cycle entrainment have been a source of significant interest in part because of the apparent disconnect between after-effects of T cycle (where *ex vivo* SCN and behavioral measurements negatively correlate) and of photoperiod (where *ex vivo* SCN and behavioral measurements agree). In our combined T cycle/photoperiod paradigm we found a significant negative relationship between T cycle length and SCN τ length. Entrainment to short T cycles *in vivo* resulted in lengthened SCN τ *ex vivo*, and *vice versa* as previously reported (Aton et al., 2004; Azzi et al., 2017; Molyneux et al., 2008). Visually, there is an intriguing reversal in trend direction of possible photoperiod effects between T24 and the two non-24 T cycles that could indicate an interaction. In T24 cycles, the previously reported trend between long and short photoperiod (shortened, behavior-like τ after-effect for long photoperiod) is present, while in T23 and T26 cycles, the trend between long and short is inverted. This relationship results in the longest and shortest τ aligning with T23 Long and T26 Short, the same two groups highlighted above (Figure 3) as overlap groups where the effects of T cycle and photoperiod are in line with one another. However, no significant photoperiod effects or interactions were revealed by the ANOVA, or by multiple post-hoc tests for differences within the T cycle groups, so these visual trends require further testing.

### *Ex vivo* SCN after-effects have characteristics similar to behavioral after-effects

To investigate the relative influence of parametric versus non-parametric inputs on SCN after-effects *ex vivo*, we collected slices from animals entrained to skeleton long and short photoperiods (Figure 6A). Similar to *in vivo* Ψ of the behavioral rhythm to skeleton photoperiods, there was a significant effect of skeleton onset-offset interval on the Ψ of the PER2∷LUC rhythm (Figure 6B). Unlike *in vivo*, the PER2∷LUC Ψ did not have a significant effect of photoperiod completeness, although there was a trend towards a decrease in the difference in Ψ between long and short versions of skeleton photoperiods compared to the complete photoperiods. The σ_ϕ_ after-effect was significantly influenced by the onset-offset interval (Figure 6C), similar to *in vivo*. The σ_ϕ_ after-effect *ex vivo* was also significantly influenced by photoperiod completeness. However, the mean difference in SCN σ_ϕ_ between the long and short versions of the skeleton photoperiods was larger compared to that of the complete photoperiods. Though the interaction that this implies was not statistically significant, it does suggest that skeleton interval is a principal driver of σ_ϕ_ after-effects similar to *in vivo* α (Figure 4C). As with behavioral τ after-effects, we found no significant effect of onset-offset interval or photoperiod completeness on the *ex vivo* τ after-effect, again suggesting that the effects of photoperiod on τ after-effects are modest compared to T cycle entrainment.

**Figure 6.**
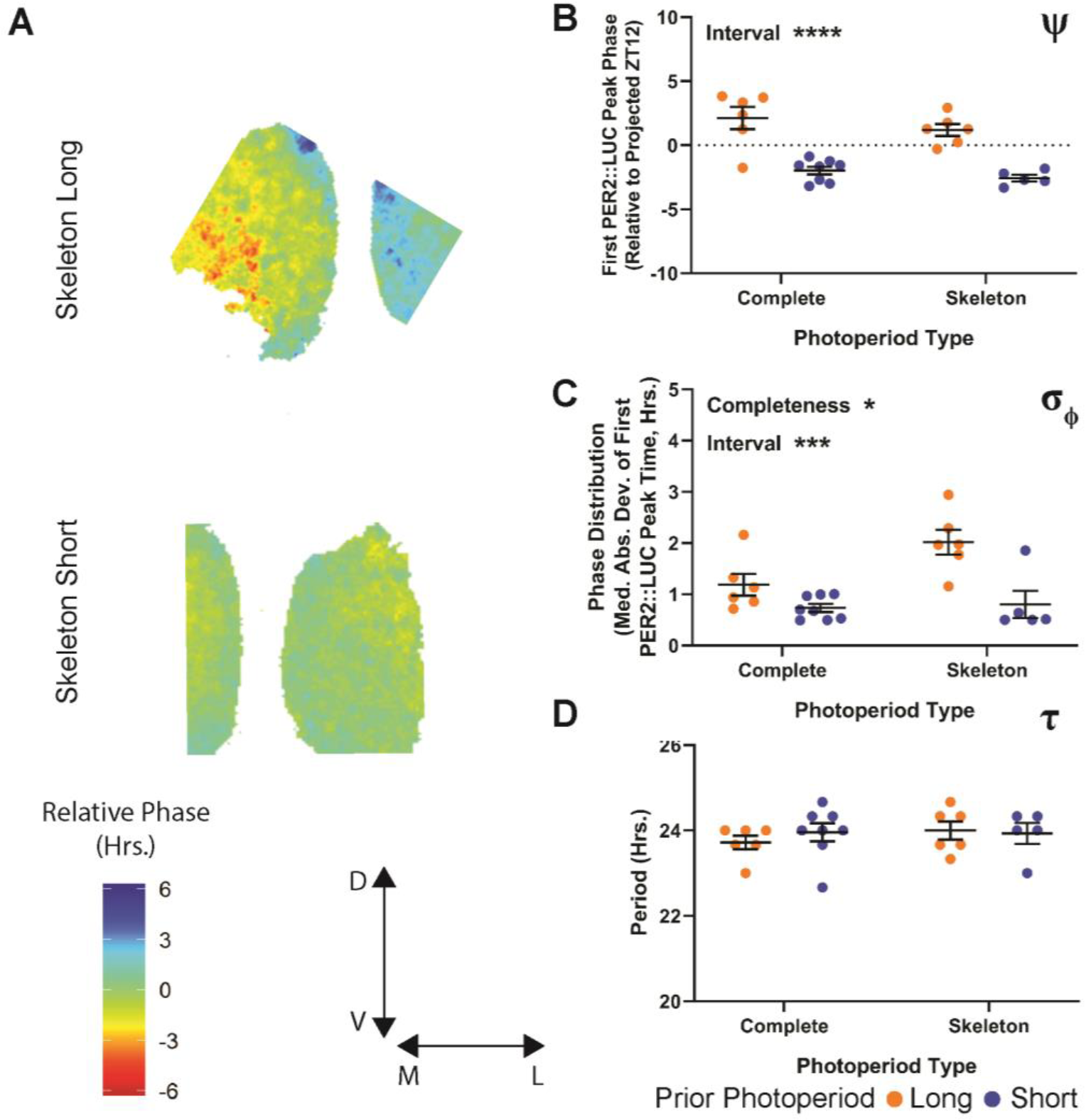
PER2∷LUC after-effects following entrainment to skeleton long and short photoperiods compared to T24 complete long and short photoperiods. **A**, representative relative phase maps from skeleton long (top) and short (bottom) photoperiod. **B**, Ψ of the PER2∷LUC rhythm following entrainment to skeleton long and short photoperiods. Values represent the time of the first PER2∷LUC peak relative to the projected lights off for the first cultured cycle (projected ZT 12). Interaction *p* = 0.7514 (0.1326%), Completeness *p* = 0.1660 (2.651%), Onset-Offset Interval *p* < 0.0001 (70.79%). **C**, σ_ϕ_ (median absolute deviation of PER2∷LUC peak times) across SCN slices from animals entrained to skeleton long and short photoperiod. Interaction *p* = 0.0654 (7.695%), Completeness *p* = 0.0318 (10.77%), Onset-Offset Interval *p* = 0.0003 (37.26%). **D**, τ of the PER2∷LUC rhythm over 5 cycles *ex vivo*. Interaction *p* = 0.4867 (2.277%), Completeness *p* = 0.5608 (1.587%), Onset-Offset Interval *p* = 0.6959 (0.7131%). Long and Short complete photoperiod values in B-D are replotted from Figure 5. Two-way ANOVA results represent *p* value and percent variation (*p* < 0.0001 ****, *p* < 0.001 ***, *p* < 0.01 **, *p* < 0.05 *). Complete Long, n = 6; Complete Short, n = 8; Skeleton Long, n =6, Skeleton Short, n = 5.

## DISCUSSION

Measurements in constant conditions reveal properties of the SCN and downstream circadian outputs free of light input, and as such the study of after-effects is one means to understand the plasticity of the SCN in response to seasonal light conditions. Understanding after-effects is also useful *per se*, as the encoding of light information within the SCN allows seasonal circadian properties to remain stable across ephemeral changes in the environment. Organisms in the wild experience fluctuations in light exposure during the annual photoperiodic cycle because of weather, nesting, hibernation, and other factors. Stable but flexible encoding of seasonal light inputs within the circadian system would allow for consistency across these light exposure fluctuations.

To improve our understanding of how after-effects are induced, we examined the relative influences of T cycle length, photoperiod, and onset-offset interval on *in vivo* (α, τ) and *ex vivo* (σ_ϕ_, τ) after-effects, as well as on Ψ (both *in vivo* and *ex vivo*). Using eight light cycle input groups, including six T cycle/photoperiod combinations and two skeleton photoperiods, we tested which specific components of light cycle input (light duration, onset-offset interval, repeated phase shifts) are encoded by the SCN in a persistent manner by measuring after-effects in circadian behavior and in SCN rhythms assayed *ex vivo*.

### Comparison of *in vivo* and *ex vivo* responses

We found that Ψ is strongly influenced by T cycle and photoperiod length, both *in vivo* (Figure 2B) and *ex vivo* (Figure 5B), with a significant interaction between T cycles and photoperiod in setting subsequent Ψ in both cases. Using T24 skeleton photoperiods to assess the influence of non-parametric (light onset-offset interval) and parametric (light interval completeness) input, we found both input types significantly influence Ψ *in vivo* (Figure 4B) while observing significant influences of non-parametric input and a trend toward an influence of parametric input *ex vivo* (Figure 6B). Determination of Ψ has long been considered to be non-parametric, with the exception of the continuity of light on full light cycles disambiguating onset from offset on near symmetrical light cycles (e.g. near T24 12:12, Pittendrigh and Minis, 1964), however our results suggest a more complex picture akin to the circadian surface entrainment model of Roenneberg for *Neurospora* (Rémi et al., 2010; Roenneberg et al., 2010). Future experiments may provide further information about the parametric and non-parametric influences of T cycles and interval lengths through the use of skeleton photoperiods with non-24 hour T cycles.

After-effects in α were strongly influenced by photoperiod length (Figure 2C) in a non-parametric manner (Figure 4C). This result was mirrored by the significant main effect of photoperiod on *ex vivo* SCN σ_ϕ_ (Figure 5C), which was also primarily non-parametric (Figure 6C). One notable difference between *in vivo* α and *ex vivo* SCN σ_ϕ_, however, was the general increase in SCN σ_ϕ_ measured in both T26 photoperiods. The increase in the T26 Short group may be related to the contracted α observed in LD during entrainment to T26 Long and Short (Figure 2A, bottom row): Though the free-running α expands on the first day of DD, there is apparent negative masking occurring in the early active phase of these mice during LD. Because slices were made during the last LD cycle rather than on the first day of DD, the σ_ϕ_ could reflect that negatively-masked pattern and imply that σ_ϕ_ is correlated with overt “net” α, rather than the underlying assumed endogenous active phase length.

We found that behavioral τ after-effects are significantly influenced by both T cycle and photoperiod length, but that the T cycle influence is of greater statistical significance and is the source of a greater percentage of the overall variation (Figure 2D), suggesting that the T cycle input to τ after-effects is more robust. Our results with T24 complete and skeleton photoperiods *in vivo* support this notion, as we did not observe significant τ after-effects with either complete or skeleton photoperiods (Figure 4D). Previous studies have described τ after-effects of complete and skeleton photoperiods on behavioral τ in mice, but with modest differences between the two skeleton photoperiod lengths (Pittendrigh and Daan, 1976a). While locomotor τ after-effects reflected the length of the entraining T cycle, we observed negative correlations between the τ after-effects of *ex vivo* SCN and the length of the entraining T cycle, consistent with previous findings (Aton et al., 2004; Azzi et al., 2017; Molyneux et al., 2008). We did not detect any statistically significant modulation of the T cycle effects on τ by varying photoperiod, but visually, there are trend reversals in the expected direction of photoperiodic effects on τ in the non-24-hour T cycle groups that suggest the possibility of interactions that we did not detect statistically (T24 Long vs. T23 Long, *p* = 0.1513; T24 Short vs. T26 Short, *p* = 0.0584; Sidak’s Multiple Comparisons, see further discussion below).

### Differences between T cycle- and photoperiod-induced responses

Because α and τ after-effects have been observed together (shorter α, shorter τ; longer α, longer τ), previous studies have attempted to explain both changes with a single underlying mechanism (Beersma et al., 2017; Gu et al., 2016). We observed different patterns of responses for after-effects in α and τ in photoperiod/T cycle combinations (Figure 2A, C-D) as well as in T24 photoperiods (Figure 4A, C-D), and correlation of changes in α and τ were specific to certain T and photoperiod combinations (Figure 3). When measuring the correspondence of τ and α after-effects *in vivo*, correlations were only observed in cases where T cycle and photoperiod effects are expected to align (Figure 3). These results suggest that while the two after-effects may frequently be observed together, their distinct patterning in response to particular T cycles and photoperiods could be driven by disparate but conditionally convergent mechanisms.

Changes in SCN σ_ϕ_ remain a likely explanation for alterations in downstream locomotor behavior duration, as an extended population-level high-firing phase of SCN neurons (for which the PER2::LUC σ_ϕ_ is a proxy here) would logically underlie the regulation of timing activity or inactivity (Ciarleglio et al., 2009, 2011; Inagaki et al., 2007; VanderLeest et al., 2007). Changes in τ, however, shown here to be induced primarily through T cycle entrainment and likely the requisite repeated phase shifts thereof, may be regulated in a different manner. Mechanisms suggested recently by other groups (Azzi et al., 2014, 2017) surrounding the epigenetic changes that may occur after extended entrainment to T cycles may help explain alterations in τ that we observed.

### Regional phase differences and their influence on after-effects

Regional phase differences within the SCN network have been observed in response to T cycles and photoperiod, and have been proposed to drive τ after-effects (Azzi et al., 2017; Buijink et al., 2016; Evans et al., 2013; Myung et al., 2015). In SCN explants from mice entrained to long photoperiods, the ventral or ventromedial region SCN phase leads in some studies (Buijink et al., 2016; Evans et al., 2013), but the reverse regional phase relationship was observed in another (Myung et al., 2015). SCN τ shortening after entrainment to long photoperiods was observed to correlate with ventral phase lead (Evans et al., 2013), with ventral phase lag (Myung et al., 2015), or not be present (Buijink et al., 2016). Similarly, Azzi *et al*. (2017) found that following entrainment to T cycles, there was an anticorrelation between SCN τ after-effect and the previous T cycle τ, as expected from previous results (Aton et al., 2004; Molyneux et al., 2008). The same study also found that shortened τ correlated with ventral phase lag, while lengthened τ correlated with ventral phase lead – the opposite pattern previously found for photoperiodic τ after-effects by the same group (Evans et al., 2013). Thus, there is no clear consensus in the literature as to whether SCN regional phase differences mediate τ after-effects.

We also observed the anticorrelation of *ex vivo* SCN τ after T cycle entrainment *in vivo*, as well as a variety of regional σ_ϕ_ across our experimental conditions that included examples of ventral core phase lead in response to short T cycles (see T23 Long in Figure 6) similar to Azzi *et al*. (2017), and examples of more ventromedial phase clustering similar to that of Buijink *et al*. (2016) in response to long T cycles (see T26 Short in Figure 6). Differences in regional patterns (ventral core vs. ventromedial) may be explained by slice position on the rostral caudal axis of the SCN. Azzi et al. (2017) show that taking sections more caudal in the SCN changes the regional phase pattern from ventral core grouping to a ventromedial grouping similar to Buijink (2016). We speculate that we may not have controlled well enough for slice depth, resulting in our observation of variations of both patterns in our data.

Conceptually, the anticorrelation of SCN *ex vivo* τ with *in vivo* τ after-effects of T cycles presents a challenge. One interpretation of these results is that the observed *ex vivo* τ reflects a deafferentation-induced reorganization of the SCN network, indicating extra-SCN input is key to T cycle after-effects *in vivo* (Aton et al., 2004; Molyneux et al., 2008). Support for this view also comes from genetic studies showing an anticorrelation between *in vivo* behavioral and *ex vivo* SCN τ effects of genetic serotonergic de-afferentation of the SCN (Ciarleglio et al., 2014), and lack of correlation of *in vivo* and *ex vivo* SCN period after-effects of genetic manipulations of SCN gene expression (Mieda et al., 2015, 2016). In addition, part of the confound regarding previous *ex vivo* SCN T-cycle after-effects appears to be a bias in the reporting of SCN regional rhythms by the standard PER2::LUC reporter that more highly weights τ of the dorsal region (Azzi et al., 2017). In any case, the lack of correspondence between *in vivo* and *ex vivo* T-cycle τ after-effects in the SCN makes it difficult to attribute *ex vivo* findings of regional patterns or mechanisms to the *in vivo* case, although the system can be used to study SCN *ex vivo* reorganization and its mechanisms (Azzi et al., 2017).

### SCN σ_ϕ_ and its relationship with α

The degree of SCN neuron σ_ϕ_ is a characteristic attribute of photoperiodic encoding, as it represents a change in state of the SCN induced by the duration of the light interval of the daily cycle and is maintained after transfer to constant conditions (Buijink et al., 2016; Ciarleglio et al., 2011; Inagaki et al., 2007; VanderLeest et al., 2007). Our results suggest that the timing of light/dark transitions is a key factor in influencing SCN σ_ϕ_ (Figure 6C) and *in vivo* α (Figure 4C), although we also detected an effect of light duration on SCN σ_ϕ_ (Figure 6C). The spread in the timing of neuronal rhythms allows for the daily duration of high firing activity of the SCN to be broadened or contracted without required alteration to the waveforms of individual SCN neurons, although this aspect is modulated by developmental inputs (Ciarleglio et al., 2011). As such, the degree of σ_ϕ_ within the SCN is a potential target for artificial SCN manipulation – driving neurons to adopt the σ_ϕ_ characteristic of a photoperiod, such as by pharmacologically, optogenetically, or chemogenetically extending the high firing phase, would be expected to induce the behavioral α characteristic of that particular photoperiod, and perhaps other photoperiod-dependent responses, such as reproductive state and affective behaviors.

Because current methods for *in vivo* imaging are limiting, the measure of α after-effects may be a useful proxy for inferring the waveform of the SCN. The integrity of this SCN waveform-α connection depends on the correlation between the two. Our findings that α and *ex vivo* SCN neuron σ_ϕ_ associate across multiple lighting conditions regardless of net daily phase shift do not address causality, but strengthen the association between the two measurements. As the broadened electrical waveform of the SCN is associated with the compressed behavioral α seen in long photoperiods (and the contracted waveform with extended α in short photoperiods), the spread in phase of SCN neurons and the corresponding widening of overall firing that occurs as a result should account for the observed changes in α (Houben et al., 2009).

Changes to SCN network synchrony may have additional ramifications in the form of altered responses to light input, potentially explaining previously observed larger phase shifts after exposure to short days, and smaller phase shifts after exposure to long days (Refinetti, 2002). The role of SCN neuron σ_ϕ_ in mediating those altered responses and altered τ has been explored computationally (Gu et al., 2016) and through altered GABA signaling (Farajnia et al., 2014). In their analysis of after-effects on τ, Pittendrigh and Daan (1976b) suggested that this change in circadian property may provide functional significance in terms of priming future responses to light, but it remains to be seen whether any of these after-effects constitutes an adaptive advantage or if they are simply a side effect of altered network synchrony.

## CONCLUSION

Taken together, our results show that distinct components of light cycles can have distinct effects on the circadian system, with (a) the timing of light/dark transitions in photoperiods driving α plasticity; (b) combined photoperiod and T cycle length driving plasticity in behavioral τ and Ψ of both *in vivo* behaviors and *ex vivo* SCN gene expression; and (c) the light/dark transition interval and long T cycle lengths affecting *ex vivo* SCN σ_ϕ_. These results open the door to selective manipulation of circadian plasticity in α or τ for further experimentation and for potential translation to treat photoperiodic syndromes. The results shown here also demonstrate the utility of deconvolving overlapping types of input when studying complex processes within the SCN. Our results indicate that α and τ after-effects are possibly driven by distinct, though perhaps related, mechanisms, and such information can be incorporated into experimental and computational approaches to test the system in the future. With new techniques currently being established in our field to measure SCN neuronal firing rate, calcium signaling, and activity levels with high cellular specificity and extended timeframe, there will be an improved ability to characterize the specific effect of photoperiodic influences on these circadian components.

## ACKNOWLEDGMENTS

The authors would like to thank Maria Luísa Jabbur, Carl Johnson, Terry Page, and Chloe Snider for their helpful comments and discussions. This work was supported by NIH R01 GM117650 to D.G.M., NIH F31 NS096813 to M.C.T., and NSF GRFP 0909667 to M.C.T.

## AUTHOR CONTRIBUTIONS

M.C.T. and D.G.M conceived the experiments. M.C.T. performed the experiments. M.C.T. and

J.J.H. analyzed the data, M.C.T., J.J.H., and D.G.M. wrote the paper.

## Supplementary Figures

**Supplementary Figure 1.**
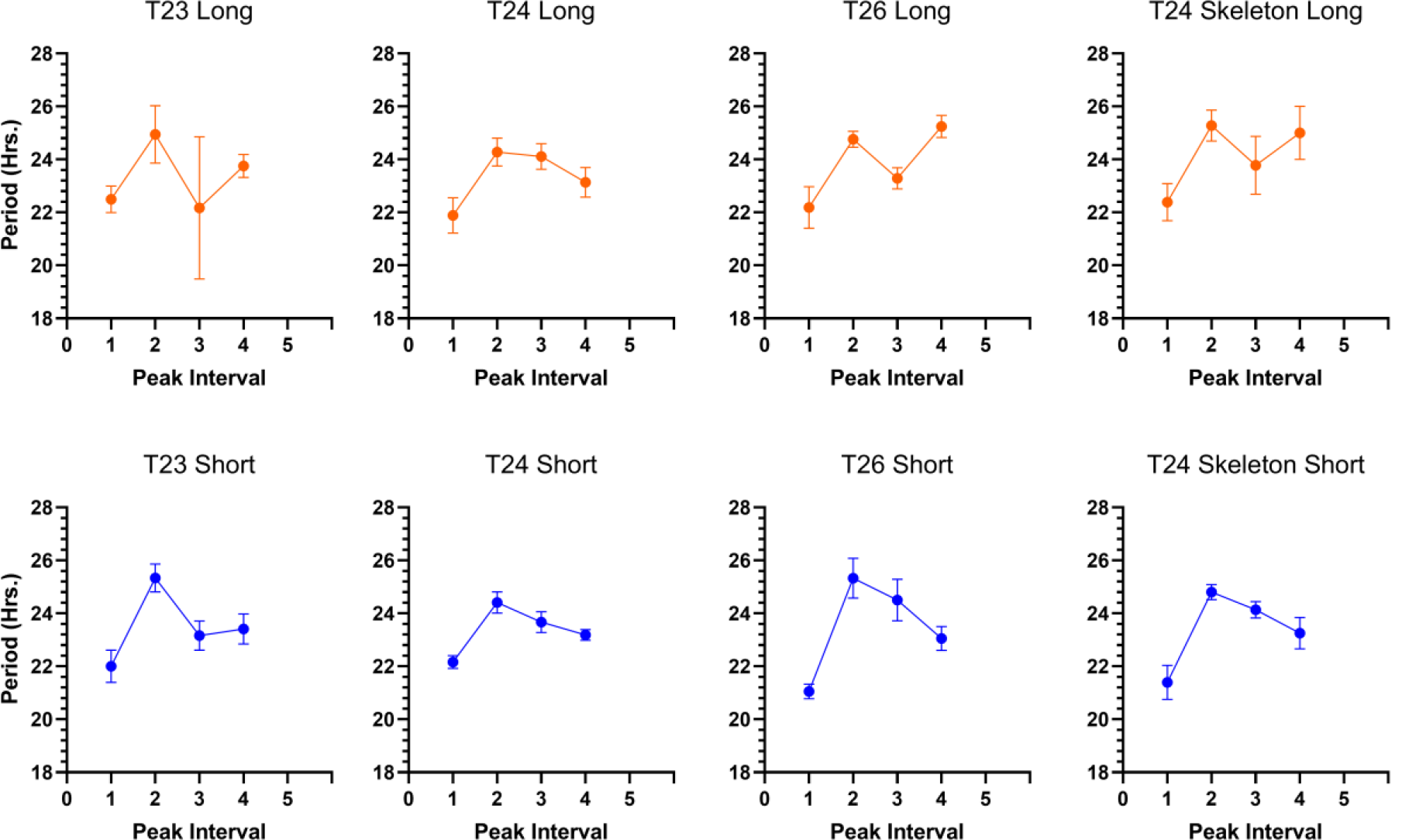
Mean period values for each interpeak interval for the first five cycles recorded *ex vivo*. Error bars represent standard error of the mean.

**Supplementary Figure 2.**
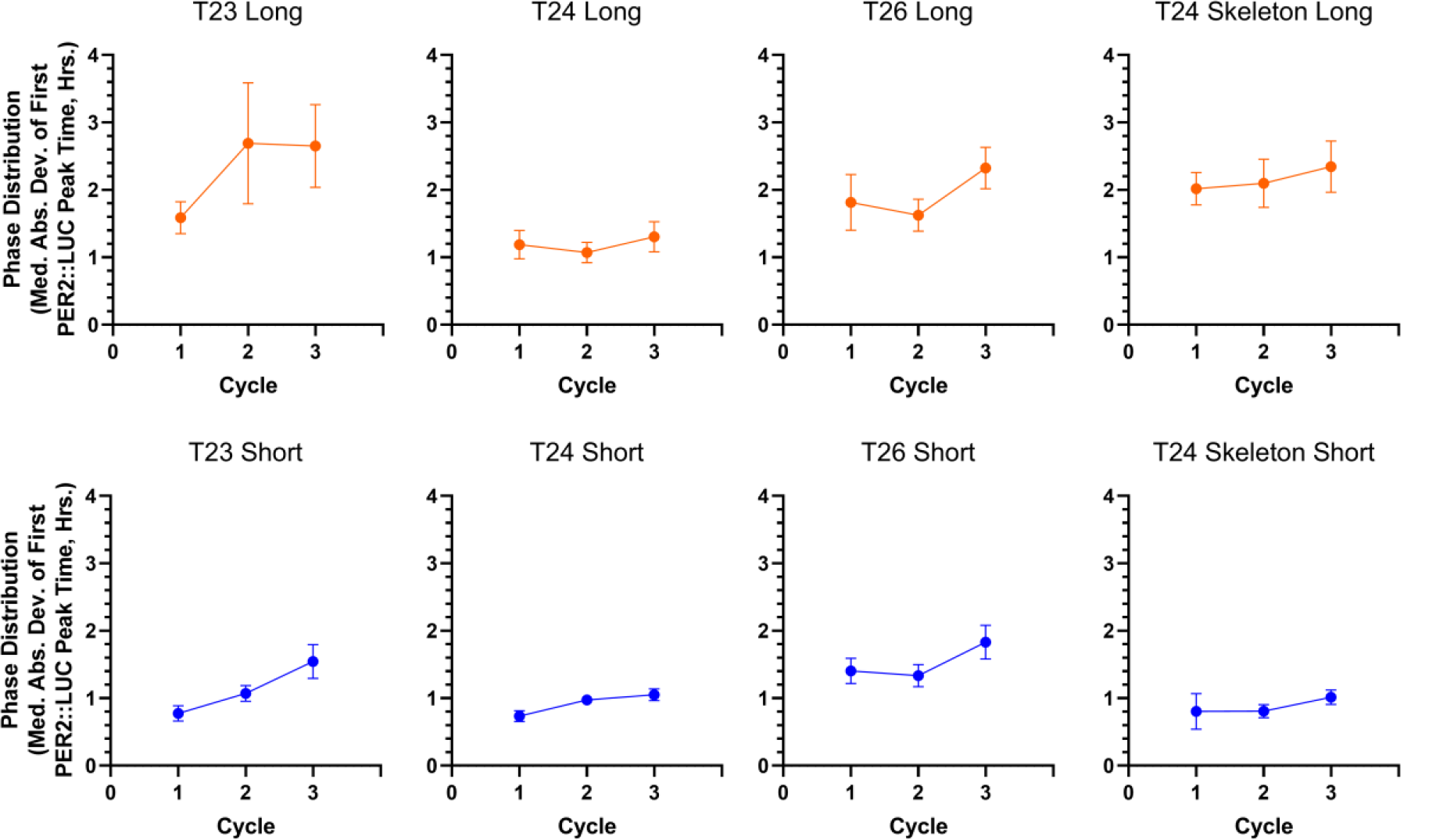
Phase distribution values calculated each cycle for the first three cycles *ex vivo*. Error bars represent standard error of the mean.

